# Cell-free expression and characterization of multivalent rhamnose-binding lectins using biolayer interferometry

**DOI:** 10.1101/2023.01.05.522863

**Authors:** Katherine F. Warfel, Eugénie Laigre, Sarah E. Sobol, Emilie Gillon, Annabelle Varrot, Olivier Renaudet, Jerome Dejeu, Michael C. Jewett, Anne Imberty

## Abstract

Lectins are important biological tools for binding glycans, but recombinant protein expression poses challenges for some lectin classes, limiting the pace of discovery and characterization. To discover and engineer lectins with new functions, workflows amenable to rapid expression and subsequent characterization are needed. Here, we present bacterial cell-free protein synthesis as a means for efficient, small-scale expression of multivalent, disulfide bond-rich, rhamnose-binding lectins. Furthermore, we demonstrate that the cell-free expressed lectins can be directly coupled with BLI analysis, either in solution or immobilized on the sensor, to measure interaction with carbohydrate ligands without purification. This workflow enables determination of lectin substrate specificity and estimation of binding affinity. Overall, we believe that this method will enable high-throughput expression, screening, and characterization of new and engineered multivalent lectins for applications in synthetic glycobiology.

## Introduction

Lectins are proteins that recognize and bind specific glycans, making them interesting scaffolds for therapeutics, diagnostics and quality control reagents for glycoprotein products (Arnaud et al. 2013; Fernandez-Poza et al. 2021). However, recombinant expression of lectins can be challenging due to properties such as toxicity, size or presence of disulfide bonds (Martínez-Alarcón et al. 2018). One strategy to overcome these challenges is cell-free expression (CFE), which harnesses biological machinery to enable high-yielding transcription and translation outside of the living cell (Carlson et al. 2012; Silverman et al. 2020). The modular and open CFE reaction environment allows for manipulation of expression conditions, enabling production of complex products including proteins containing disulfide bonds (Goerke and Swartz 2008; Dopp and Reuel 2020), membrane proteins (Matthies et al. 2011; Kruyer et al. 2021), and glycosylated proteins (Kightlinger et al. 2019; Hershewe et al. 2021; Stark et al. 2021; Jaroentomeechai et al. 2022). CFE is also scalable from the nanoliter to liter scale (Zawada et al. 2011), allowing for small-scale parallel expression of many proteins simply by switching out the template DNA added to the reaction, and accelerating protein screening (Hunt et al. 2021; Hunt et al. 2022).

To take advantage of CFE for rapid lectin screening, it would be ideal to assess the functionality of produced lectins directly in the reaction without purification. Biolayer interferometry (BLI) is a technique that has been recently used to characterize protein binding parameters without a purification step, indicating possible compatibility with CFE reactions (Pogoutse et al. 2016). In addition, BLI has been used to study multivalent lectin-carbohydrate interactions in purified systems (Laigre et al. 2018; Picault et al. 2022). Notably, this technique requires less material than surface plasmon resonance (SPR) or isothermal titration calorimetry (ITC), is label free, and can be run in a plate-based format, making it more compatible with small-scale cell-free expression (Concepcion et al. 2009).

To demonstrate our cell-free workflow, we selected two model eukaryotic lectins, SUL-I (PDB-ID 5H4S) from *Toxopneustes pileolus* (sea urchin) venom and CSL3 (PDB-ID 2ZX2) from *Oncorhynchus keta* (chum salmon) eggs, that are characterized, but are difficult to isolate naturally or to produce in reasonable quantities in *Escherichia coli* (Shiina et al. 2002; Hatakeyama et al. 2015). These dimeric lectins are of interest due to their binding affinity to the glycosphingolipid globotriaosylceramide (Gb3), a tumor associated glycolipid, as well as to rhamnose-containing bacterial O-antigens (Shiina et al. 2002; Siukstaite et al. 2021). Here, we show that these rhamnose-binding lectins are soluble and active when expressed in a bacterial cell-free system. Further, we show that cell-free expressed lectins are compatible with BLI binding assays in crude reactions without purification, with lectins either immobilized or in solution. We believe that this demonstration will enable future high-throughput screening and characterization of the specificity and affinity of predicted and engineered lectins.

## Results

### Cell-free expressed multivalent lectins are compatible with direct in-solution BLI analysis

Each SUL-I monomer presents 3 binding sites with 13 disulfide bonds, while each CSL3 monomer has 2 binding sites with 8 disulfide bonds and both of them are dimeric in solution (**Figure S1A-B**) (Shirai et al. 2009; Hatakeyama et al. 2017). To support proper protein folding, SUL-I and CSL3 were produced in an *E. coli* cell-free expression system using an extract derived from T7 SHuffle cells (Dopp and Reuel 2020) and additives optimized for disulfide bond expression (**Figure 1A, Table S1**). We produced 8.2 *μ*M (274 *μ*g/mL) and 3.9 *μ*M (94 *μ*g/mL) full-length, soluble SUL-I and CSL3 respectively, as determined by radiolabeled amino acid (^14^C-Leucine) incorporation and quantification (**Figure 1B, S2A-C**). Importantly, lowering the expression temperature to 16 °C was necessary to increase the solubility and yields of both lectins (**Figure S2A-B**).

**Figure 1.**
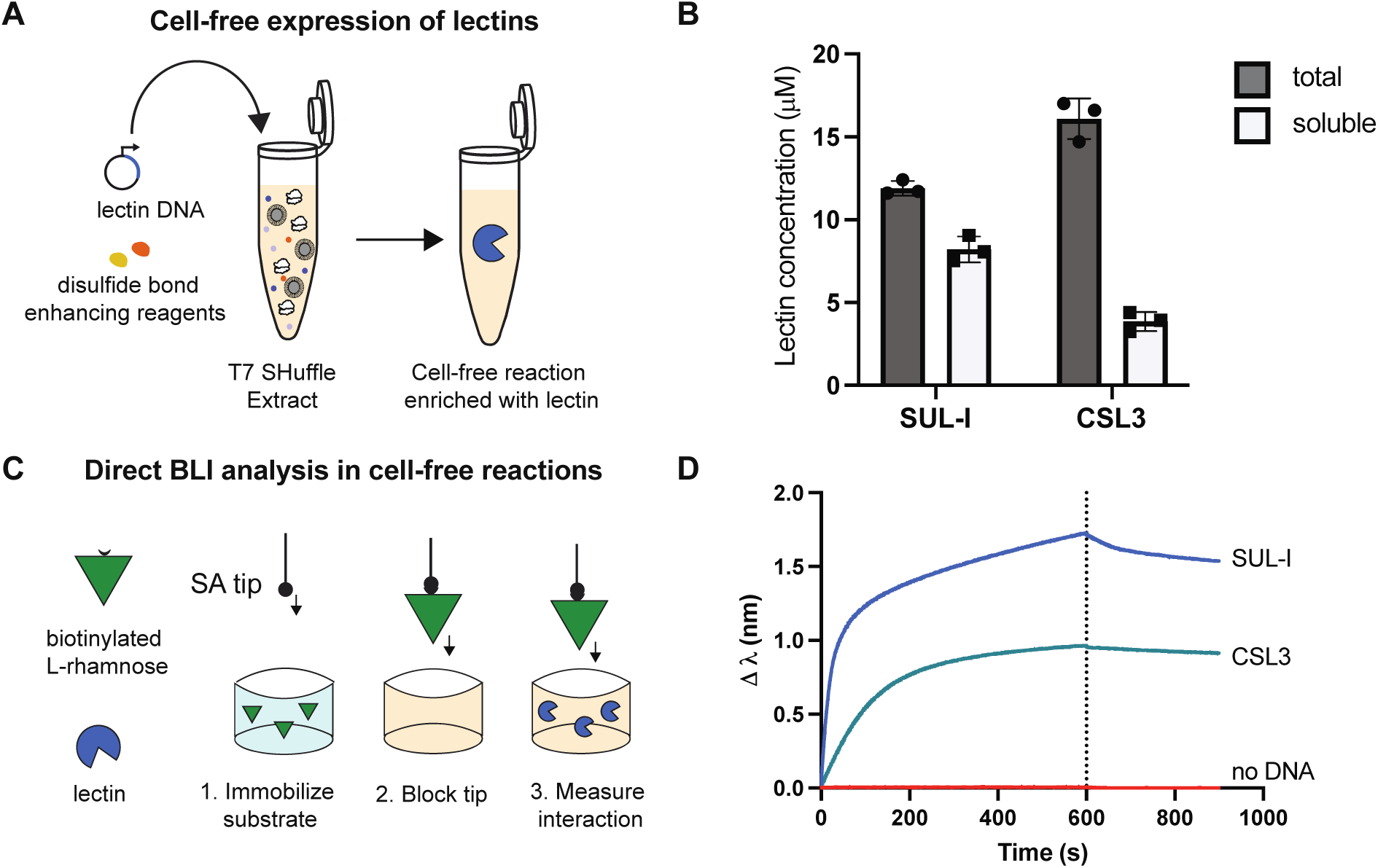
Cell-free expressed rhamnose-binding lectins are directly compatible with BLI analysis. A) Schematic representation of cell-free expression system for disulfide bond containing lectins. B) Average total and soluble yields of SUL-I and CSL3 expressed in cell-free at 16°C by radiolabeled amino acid (^14^C-Leucine) incorporation. Error bars represent the standard deviation for n=3 CFE reactions. C) Schematic of BLI assay where a biotinylated rhamnose monosaccharide is immobilized on the tip, followed by blocking with a cell-free reaction with no DNA template, and then measuring the interaction with a lectin synthesized in the cell-free reaction. D) BLI sensorgram of the interaction between a rhamnose monosaccharide immobilized on the tip and a 10-fold dilution in PBS of either negative control (no DNA template), SUL-I (∼800 nM), or CSL3 (∼400 nM) enriched cell-free reaction.

We first confirmed that cell-free expressed SUL-I and CSL3 were active by isolating them using either D-lactose or D-galactose affinity resin respectively (**Figure S2D**). Next, we determined if BLI analysis was directly compatible with unpurified lectins in cell-free reactions. We loaded biotinylated L-rhamnose monosaccharide (L-Rha*α*-sp3-biot) onto a streptavidin BLI tip. We then accounted for background interaction from the cell-free reaction mixture by adding a “blocking” step in a negative cell-free reaction (containing water instead of plasmid DNA encoding the lectin) following rhamnose substrate immobilization on the tip and prior to the interaction assay (**Figure 1C, Figure S3**). We observe a rapid and strong association phase for both lectins, with almost no dissociation phase, when cell-free reactions expressing either SUL-I or CSL3 are present in solution (**Figure 1D**). No interaction occurs in reactions that did not contain lectin DNA expression template (no DNA), confirming the specificity of the method (**Figure 1D**). The signal variation was different for the two lectins at increased dilution, indicating better binding from SUL-I. This may be related to the higher expression levels for SUL-I in the cell-free reaction and by the larger number of binding sites available in SUL-I compared to CSL3 (**Figure S4**). For SUL-I, the signal level remains unchanged with a 50-fold dilution, indicating that the dissociation constant (k_D_) is lower than the lectin concentration at this dilution (160 nM). Whereas for CSL3, we observe a more rapid decrease of the signal with increasing dilution (80 nM), yielding an approximation of k_D_ that is higher than the concentration of CSL3 at this dilution (between 80 and 400 nM).

### Cell-free expression coupled with BLI enables determination of SUL-I binding specificity and affinity

To further validate this experimental set up, we tested the specificity of SUL-I for different rhamnose substrate architectures immobilized on the BLI tip (**Figure 2A, Figure S4**). In addition to the monosaccharide, we also assessed two synthesized multivalent compounds, a tetravalent (Rha4) and a hexadecavalent (Rha16) cyclopeptide functionalized with L-rhamnose by triazole linkages, using architectures that have previously been used to assay multivalent lectin/substrate interactions (Picault et al. 2022). We also examined a commercial multivalent polyacrylamide polymer functionalized with rhamnose (PAA-rhamnose). We observed higher signal as the valency of the substrate on the tip increased, with the smallest change in wavelength for the monosaccharide and the largest change in wavelength (∼ 8-fold higher) for the Rha16 substrate, indicating that more SUL-I was able to bind to the additional L-rhamnose immobilized on the tip (**Figure 2A**). Very little dissociation was observed in all cases.

**Figure 2.**
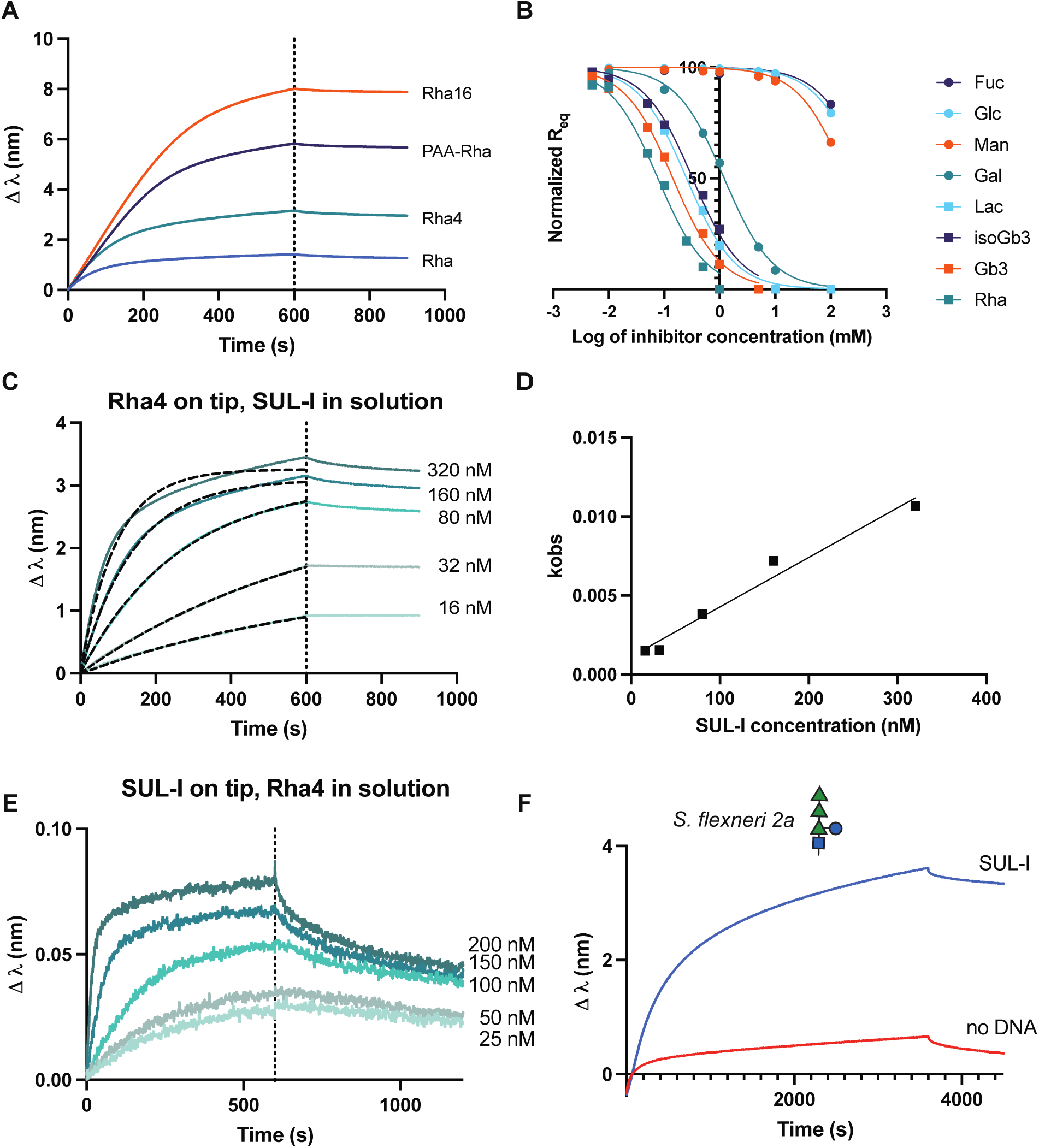
Cell-free expression coupled with biolayer interferometry enables determination of lectin binding specificity and affinity. A) Sensorgram of the interaction between different rhamnose substrate architectures (Rha16, PAA-Rha, Rha4, Rha monosaccharide) immobilized on the SA BLI tip with 50-fold diluted SUL-I (∼160 nM) cell-free reaction in solution. B) Impact of different sugar inhibitors on the interaction between the rhamnose monosaccharide tip and SUL-I lectin in solution. The normalized equilibrium binding signal (R_eq_) as determined by a 1:1 association fit, is plotted as a function of the log (inhibitor concentration) for each sugar inhibitor. Nonlinear curve fit using Graphpad PRISM log(inhibitor) vs normalized response is shown in the corresponding color for each inhibitor. C) Interaction of Rha4 substrate immobilized on the SA tip, with varying dilutions of cell-free reaction containing SUL-I (∼16 nM - 320 nM) in solution. The 1:1: model fit used to determine k_obs_ is shown in a black dotted line for each concentration and estimated concentration of SUL-I in solution is indicated to the right of the corresponding curve. D) k_obs_ values plotted as a function of SUL-I concentration and fit with a simple linear regression in Graphpad PRISM for Rha4. E) Interaction of SUL-I substrate immobilized on the Ni-NTA tip from a cell-free reaction, with varying dilutions of Rha4 substrate (∼25nM - 200 nM) in solution. F) Detection of 100-fold dilution of crude extract enriched with *Shigella flexneri 2a* O-antigen in solution with SUL-I in cell-free reaction (blue) or control cell-free reaction with no DNA (red) immobilized on the Ni-NTA tip. The monomer structure of the O-antigen glycan is indicated above the sensorgram. All data is representative of at least two independent experiments.

Additionally, we employed this strategy to determine the specificity of SUL-I by screening the ability of different mono- and oligo-saccharides (rhamnose, Gb3, isoGb3, lactose, galactose, glucose, fucose, and mannose) to inhibit the binding interaction with the rhamnose monosaccharide on the tip. We determined that a 50-fold dilution of SUL-I CFPS (∼ 160 nM) still resulted in robust signal (**Figure S4**) and incubated the diluted cell-free reaction with sugar competitors at varying concentrations for one hour before the interaction step. For all experiments where the sugar acted as an inhibitor, the signal decreased with increased concentration of inhibitor (**Figure S5**). Subsequently, we compared the equilibrium response R_eq_ (maximum equilibrium change in wavelength), as determined by fitting a 1:1 association curve for each condition, to estimate relative IC50 values for each sugar (**Figure 2B, Figure S5**) (Orthwein et al. 2021). Rhamnose had the lowest IC50 for this system (0.07 mM), followed by Gb3 (0.15 mM), while galactose had the highest observed IC50 (1.26 mM) (**Figure 2B, Figure S5A, Table S2**). Fucose, glucose, and mannose did not fully inhibit binding at the concentrations tested (**Figure 2B**).

Finally, we demonstrated the utility of this method for estimating binding kinetics and affinity (**Figure 2C**). As previously described for multivalent lectin-ligand systems with weak dissociation, we estimated the kinetic parameters k_on_, k_off_, and k_D_ from the linear relationship between k_obs_, fit by a 1:1 association model, and the lectin concentration, determined by ^14^C-Leucine incorporation (**Figure 2C-D, Figure S6**) (Picault et al. 2022). Values of k_on_ were not affected much by the architecture of the substrates, with variations from 3.5 × 10^−5^ M^-1^ s^-1^ to 1.9 × 10^−5^ M^-1^ s^-1^ (**Table S3**). We obtained the highest k_D_ of ∼ 80 nM for the monosaccharide and slightly higher affinity k_D_ (∼ 30-40 nM) for the Rha4, Rha16 and PAA-Rha on the tip (**Figure S6, Table S3**). When the rhamnose ligands are immobilized, all sensors present rhamnose multivalently regardless of individual ligand structure, resulting in little difference in binding behavior between the architectures.

We also reversed the experiment and immobilized the lectins with N-terminal 6xHis tag from a crude cell-free reaction using Ni-NTA tips, to measure interaction with rhamnose ligands in solution (**Figure 2E, Figure S7A**). This system did not follow the same kinetics as when the rhamnose ligand was on the tip, possibly due to different affinity when the lectin is immobilized, preventing a 1:1 binding analysis. However, using steady state analysis we were able to estimate a k_D_ of ∼ 100 nM for Rha4 and ∼ 50 nM for Rha16 with SUL-I on the tip (**Figure S8, Table S4**). Importantly, these k_D_ values are the same order of magnitude as our estimates for SUL-I binding affinity when the rhamnose substrate was immobilized, although the Rha16 has higher affinity than Rha4 when SUL-I is immobilized.

While we could detect interaction with the Rha4 and Rha16 ligands in solution, this experimental set up is limited by the detectable ligand size, and we could not detect binding with the rhamnose monosaccharide in solution (**Figure S7B**) as previously observed for other monosaccharides (Laigre et al. 2018). However, this method is also of interest for detection of glycans in crude samples. Towards this goal, we show that SUL-I enables detection of the *Shigella flexneri* O-antigen glycan, which contains terminal rhamnose residues (Anderson et al. 2016), in crude cell-free glycoprotein synthesis (CFGpS) extracts (Jaroentomeechai et al. 2018; Kightlinger et al. 2019; Warfel et al. 2022) without purification (**Figure 2F**). In these experiments, we accounted for background binding to the Ni-NTA tip with a reference sensor that was loaded with cell-free reaction that did not contain the DNA expression template for the lectin (**Figure S7, S8**), shown without subtraction in **Figure 2F**. Nevertheless, as previously described for protein loading onto BLI tips from crude cell lysate, a stronger, more specific interaction such as biotin-streptavidin may result in more uniform and saturated binding of the lectin to the tip (Pogoutse et al. 2016).

## Discussion

This work couples a bacterial cell-free expression platform directly with biolayer interferometry (BLI) to create a workflow compatible with small-scale expression and screening of lectins in high-throughput. As a proof of concept, we expressed two multivalent rhamnose-binding lectins from marine organisms and demonstrated that lectins in crude cell-free reactions are compatible with BLI interaction assays when tips are functionalized with glycan ligands. Importantly, we demonstrated this method with two commercially available biotinylated rhamnose ligands as well as two synthesized rhamnose substrates of varying sizes, enabling wide use of this technique. This approach does not require immobilization of the lectin and enables in-solution determination of binding specificity as well as affinity. Further, the ability to quantify the cell-free expressed protein in the crude mixture enables kinetics measurements using this method.

We also demonstrated that SUL-I expressed in a crude cell-free reaction can be immobilized on the BLI tip and used for qualitative detection of glycan epitopes in solution. While immobilization of the lectin has some drawbacks, such as size of glycoconjugate that can be detected or the impact of immobilization on binding activity, it can enable detection of glycan substrates from crude mixtures. Here, we demonstrate detection of the *S. flexneri* 2a O-antigen, of interest for vaccine development, directly in cell extract, aligning with previous reports that rhamnose binding lectins can bind smooth LPS from other Shigella serotypes (Shiina et al. 2002).

In total, we have shown that unpurified lectins in cell-free protein synthesis reactions can be used directly in BLI experiments, either in solution or immobilized on the sensor tip, for qualitative detection of carbohydrate-lectin interactions. This work advances efforts to characterize binding interactions without a purification step, which could expedite discovery and screening of new binders (Khavrutskii et al. 2013; Pogoutse et al. 2016). We expect that this workflow can be coupled to tools like LectomeExplore (Bonnardel et al. 2021) to express and screen uncharacterized lectins, and will advance the field of synthetic glycobiology for lectin discovery and application.

## Materials and Methods

### Cell-free Protein Synthesis (CFPS)

Cell-free protein synthesis and ^14^C-Leucine quantification were performed as described previously using a modified PANOx-SP (PEP) formulation in an extract prepared from SHuffle T7 Express (NEB) cells (Warfel et al. 2022). Extract was pre-incubated with 50 *μ*M IAM at room temperature for 30 minutes. Reactions were supplemented with 10 *μ*M DsbC, 1 mM GSH, and 4 mM GSSG and incubated at 16 °C for 20 hours for lectin expression. Detailed methods can be found in the Supplementary Information.

### Bio-layer interferometry (BLI)

All BLI experiments were performed using an Octet RED96 Instrument with data collected with ForteBio DataAcquisition9, analyzed and fit with ForteBio DataAnalysis9, and plotted with Graphpad PRISM. Detailed methods can be found in the Supplementary Information.

## Acknowledgements

The authors would like to thank Derek Wong, Dr. Simona Notova, Maddie DeWinter, and Dr. Andrew Hunt for helpful scientific discussions, and Dr. Ashty Karim for help with manuscript preparation. The authors also thank Hugues Bonnet for assistance with the BLI equipment. K.F.W. thanks the Chateaubriand Fellowship program: this material is based upon research supported by the Chateaubriand Fellowship of the Office for Science & Technology of the Embassy of France in the United States. A.I. acknowledges support from Glyco@Alps (ANR-15-IDEX-0002) and Labex Arcane/CBH-EUR-GS (ANR-17-EURE-0003). M.C.J. acknowledges DTRA (HDTRA1-21-1-0038) and the National Science Foundation (MCB 1936789). The authors also acknowledge support from ICMG UAR 2607 for BLI facilities.

