## Supplementary Information for "Cell-free expression and characterization of multivalent rhamnose-binding lectins using biolayer interferometry"

Supplementary Figures 1-8
Supplementary Tables 1-4
Supplementary Methods
Supplementary References

### Supplementary Figures

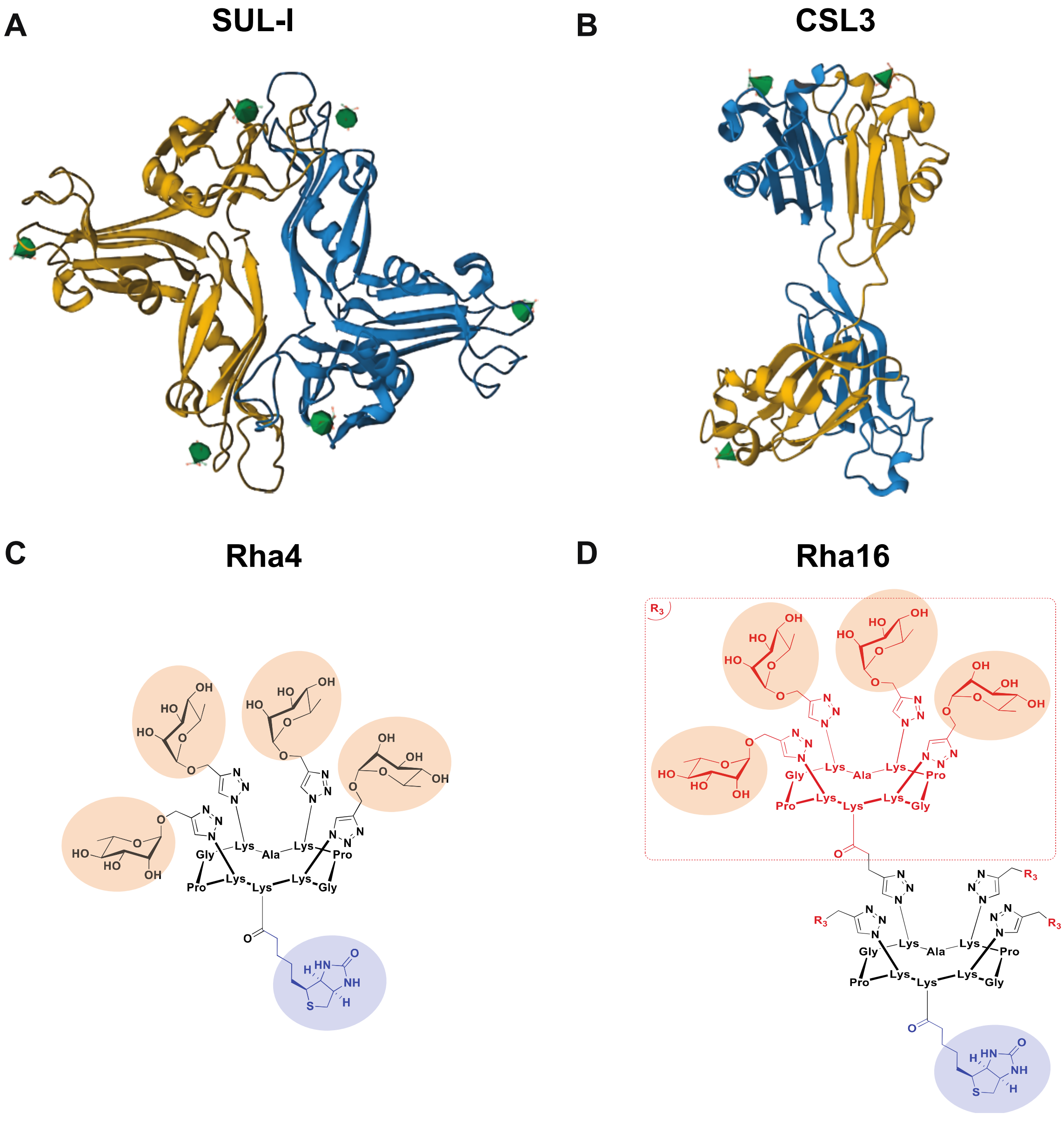


**Figure S1. Structures of SUL-I and CSL3 rhamnose binding lectins and synthesized rhamnose substrates Rha4 and Rha16.** Representation of the overall structures of (A) SUL-I (PDBID 5H4S) and (B) CSL3 (PDB-ID 2ZX2). Dimers are displayed with one protomer in gold and the other protomer in blue. Rhamnose ligands bound in structure are shown with green triangles. Structures of the synthesized ligands (C) tetravalent Rha4 and (D) hexadecavalent Rha16. The cyclopeptides are functionalized with L-rhamnose by triazole linkages.

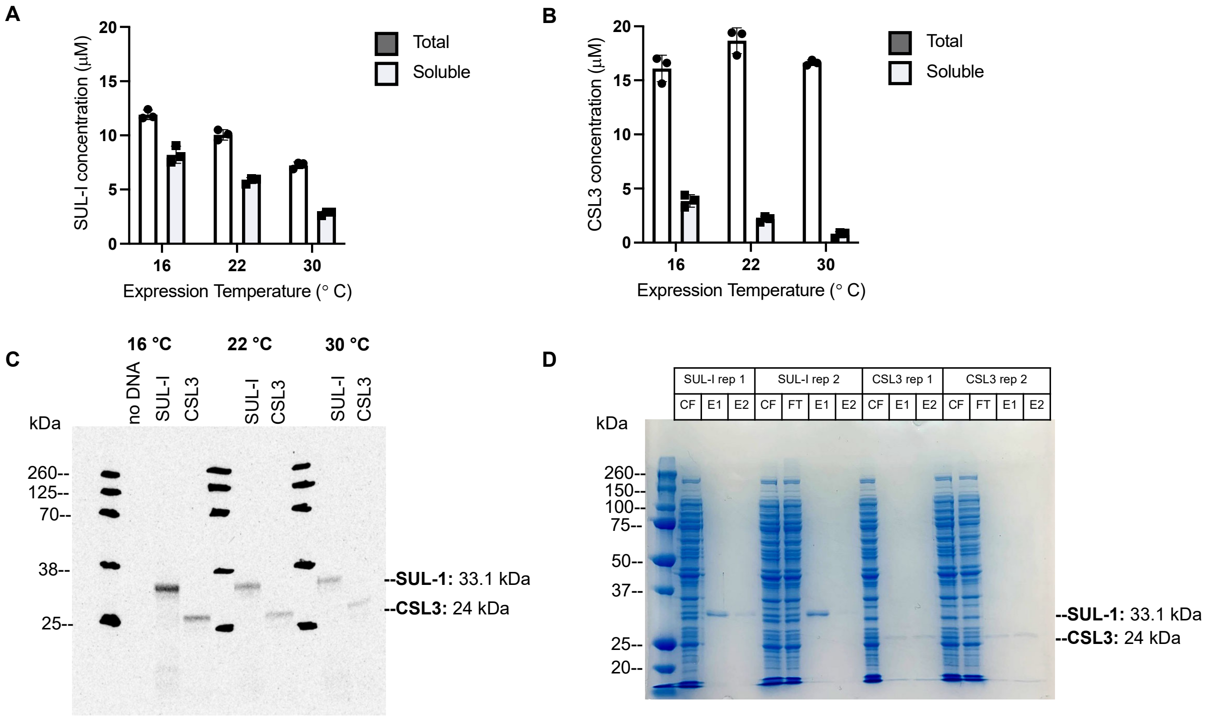


**Figure S2. Rhamnose binding lectins SUL-I and CSL3 are soluble and active when expressed using cell-free protein synthesis.** Concentration ($\mu$M) of total and soluble SUL-I (A) and CSL3 (B) expressed in cell-free reactions at three different temperatures as determined by $^{14}$C-Leucine incorporation and quantification. Error bars represent the standard deviation for n=3 cell-free protein synthesis reactions (C) Autoradiogram of the soluble fraction of cell-free reactions quantified in A-B showing protein produced with either no DNA (negative control), SUL-I, or CSL3 DNA. D) Results of affinity purification of SUL-I with lactose resin and CSL3 with galactose resin for two replicates of each purification. Soluble cell-free reaction (CF), elution fraction 1 (E1), and elution fraction 2 (E2) are shown for both replicates 1 and 2. Column flow through (FT) after binding of the reaction to the resin is also shown for replicate 2 of each purification.

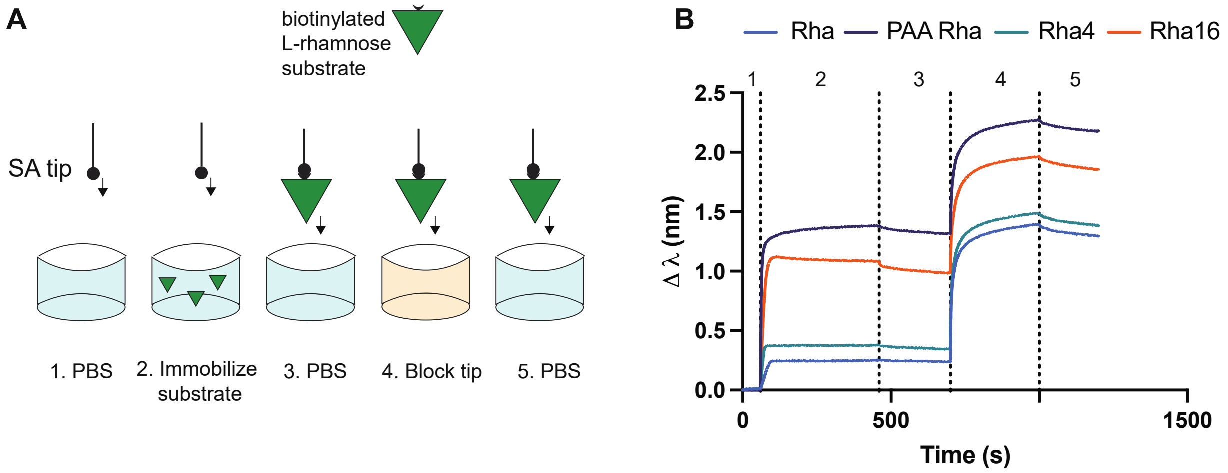


**Figure S3. Substrate immobilization and blocking procedure for streptavidin tips.** (A) Schematic illustrating the different steps of biotinylated substrate immobilization and loading to the streptavidin BLI tip. (B) Sensorgram showing the change in wavelength for tips loaded with each rhamnose substrate. Loading profile is representative of at least 3 independent experiments.

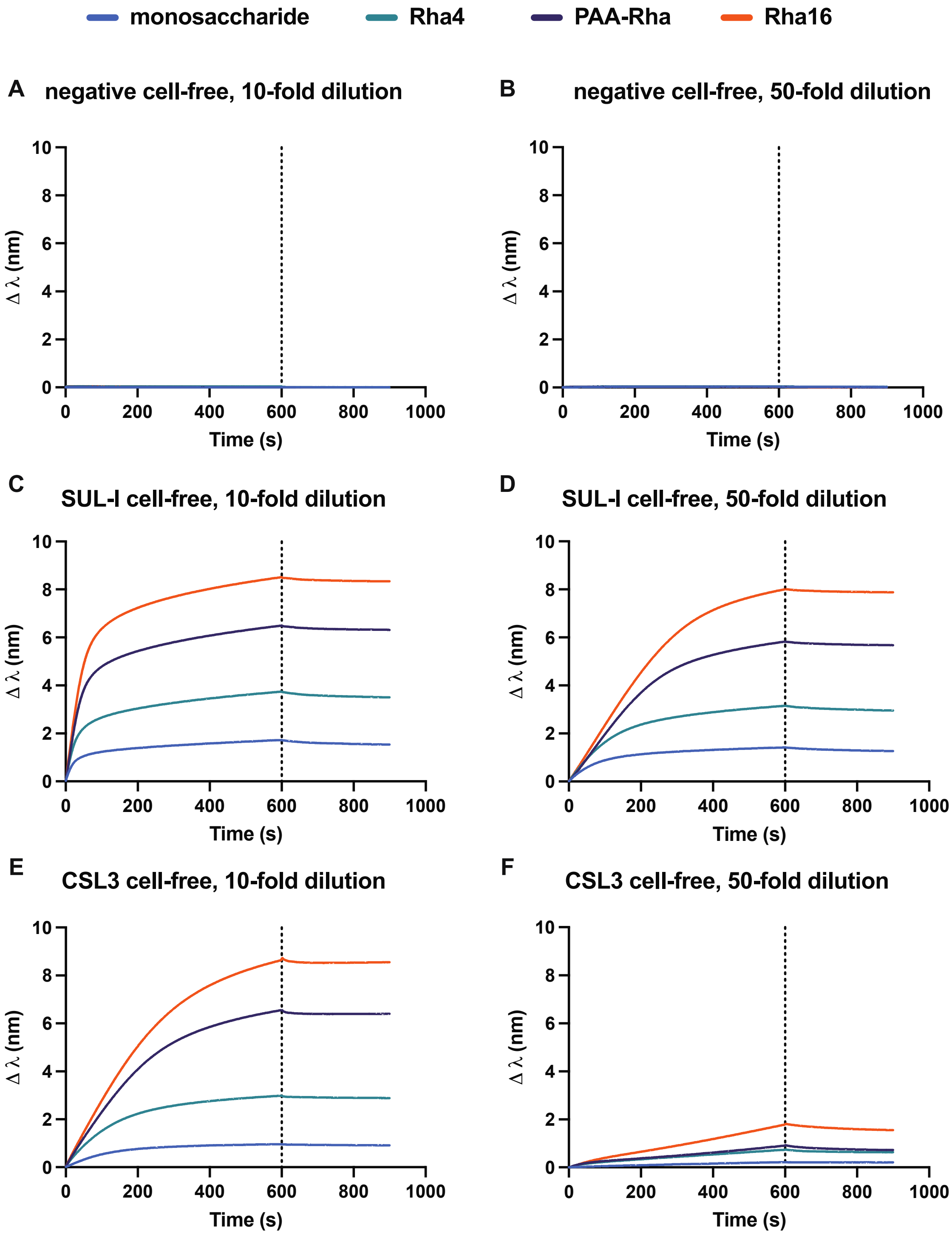


**Figure S4. Interaction between immobilized rhamnose substrates on tip with negative control cell-free reaction, SUL-I, and CSL3 at two different dilutions in PBS.** Sensorgrams showing the interaction between L-rhamnose monosaccharide, Rha4, PAA-Rha, and Rha16 immobilized on tips with a (A) 10 fold-dilution of negative cell-free reaction, (B) 50 fold-dilution of negative cell-free reaction, (C) 10 fold-dilution of SUL-I cell-free reaction, (D) 50 fold-dilution of SUL-I cell-free reaction, (E) 10 fold-dilution of CSL3 cell-free reaction, (F) 50 fold-dilution of CSL3 cell-free reaction in solution. Curves are representative of 3 independent experiments.

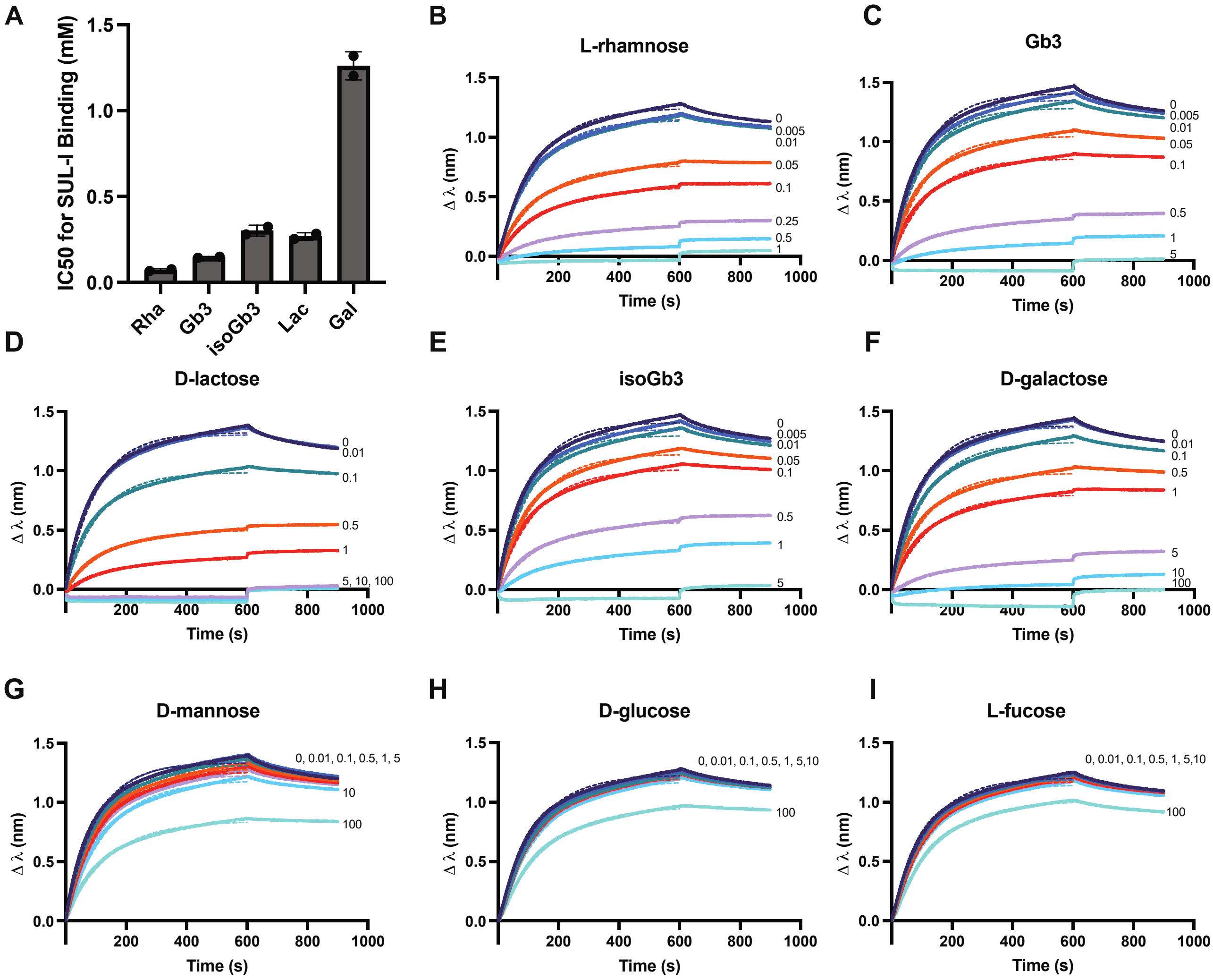


**Figure S5. IC50 values can be determined from interaction of rhamnose monosaccharide on SA tip and cell-free SUL-I pre-incubated with inhibitor in solution.** (A) IC50 of different mono- or oligo-saccharides determined by inhibition of interaction between rhamnose monosaccharide on the tip and 50-fold diluted SUL-I (~ 160 nM) cell-free reaction in solution. BLI sensorgrams resulting from interaction of rhamnose monosaccharide on the tip with SUL-I in solution incubated with varying concentrations of (B) L-rhamnose, (C) Gb3, (D) D-lactose, (E) isoGb3, (F) D-galactose, (G) D-mannose, (H) D-glucose, and (I) L-fucose. The concentration of inhibitor (mM) in solution is indicated on the right of the corresponding curve. The 1:1 model fit used to determine $R_{eq}$ is shown in a dotted line of the same color as the corresponding data. This figure highlights one replicate used in **Figure 2A**, and results are representative of two independent experiments.

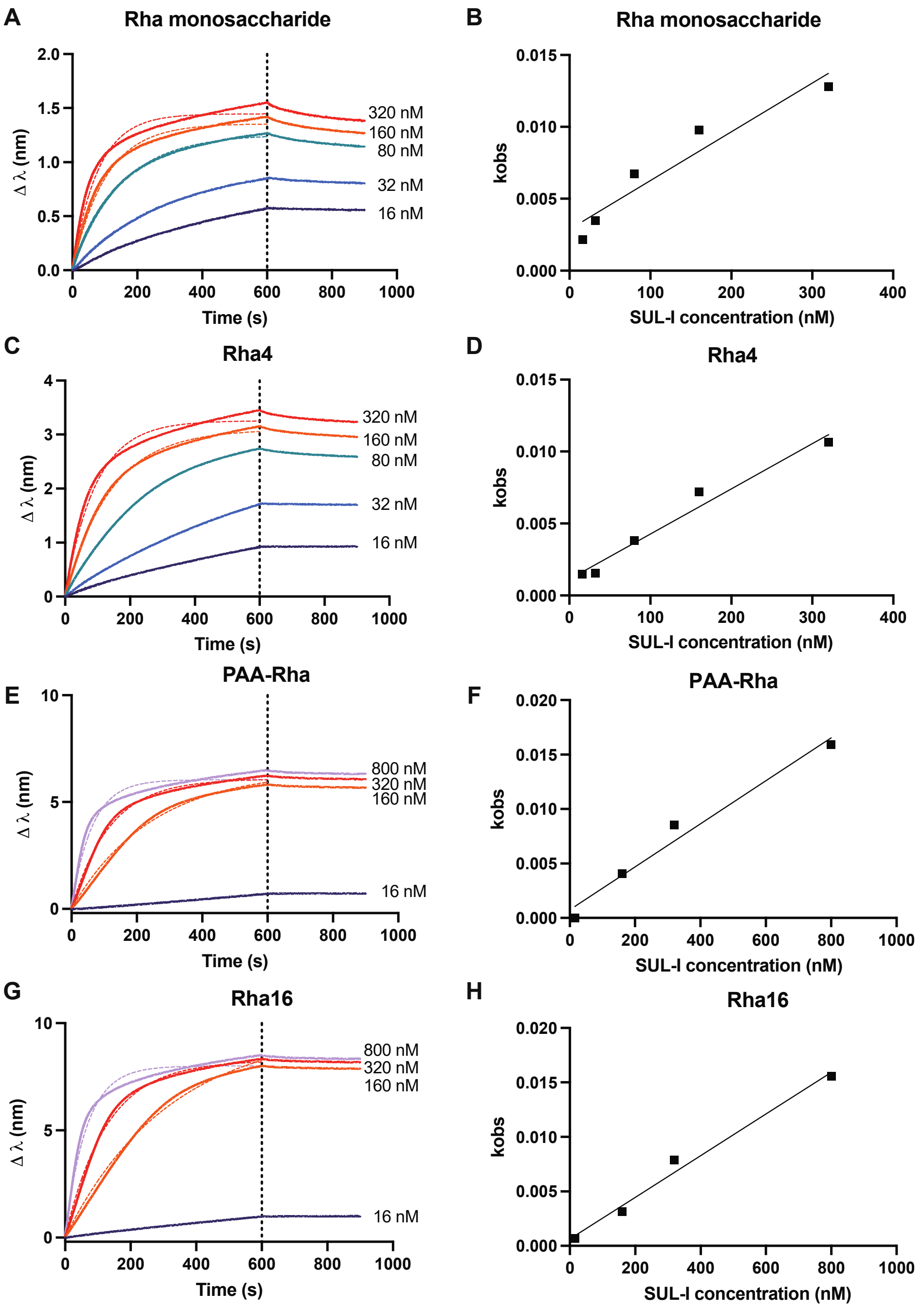


**Figure S6. Estimation of kinetic parameters with cell-free expressed lectin in solution and rhamnose substrate immobilized on tip.** Dilutions of cell-free SUL-I in solution with either (A) Rha monosaccharide (C) Rha4, (E) PAA-Rha, or (G) Rha16 immobilized on the tip. The estimated concentration of SUL-I in solution is indicated to the right of the corresponding curve. The 1:1 model fit used to determine $k_{obs}$ is shown in a dotted line of the same color as the corresponding data. The $k_{obs}$ values are plotted as a function of SUL-I concentration and fit with a simple linear regression in Graphpad PRISM for (B) Rha monosaccharide (D) Rha4, (F) PAA-Rha, or (H) Rha16 immobilized on the tip. This figure highlights one replicate used in **Figure 2C**, and results are representative of two independent experiments. Fit and corresponding kinetic parameters from this data are reported in **Table S1** as replicate 1.

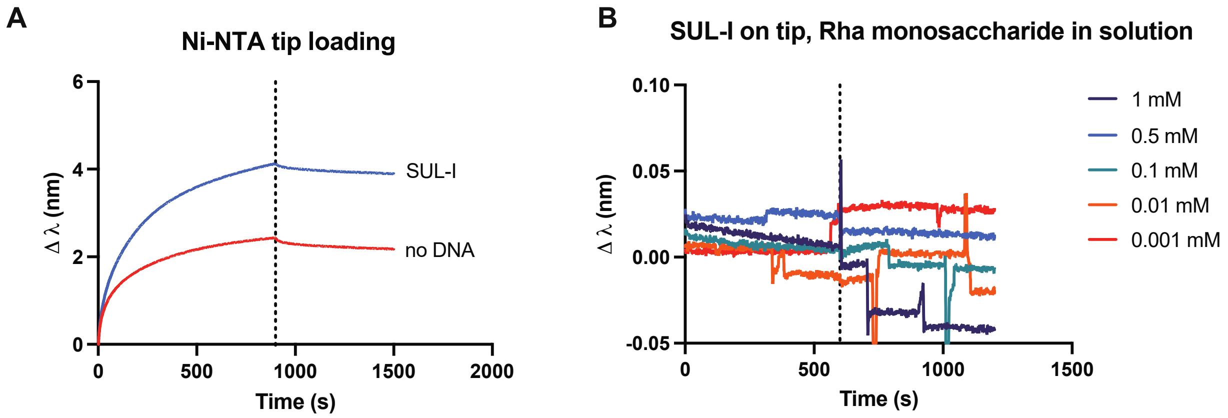


**Figure S7. His-tagged lectin can be immobilized on Ni-NTA tips from crude mixture.** (A) Change in wavelength corresponding to the loading step for either negative control (no DNA) cell-free reaction or SUL-I cell-free reaction onto a Ni-NTA tip. Non-specific binding from components of the cell-free reaction to the Ni-NTA tip is observed in the no DNA condition. (B) No interaction is observed when SUL-I is immobilized on the tip and rhamnose monosaccharide is in solution due to the size of the monosaccharide. The "no DNA" sensor is subtracted as the reference sensor.

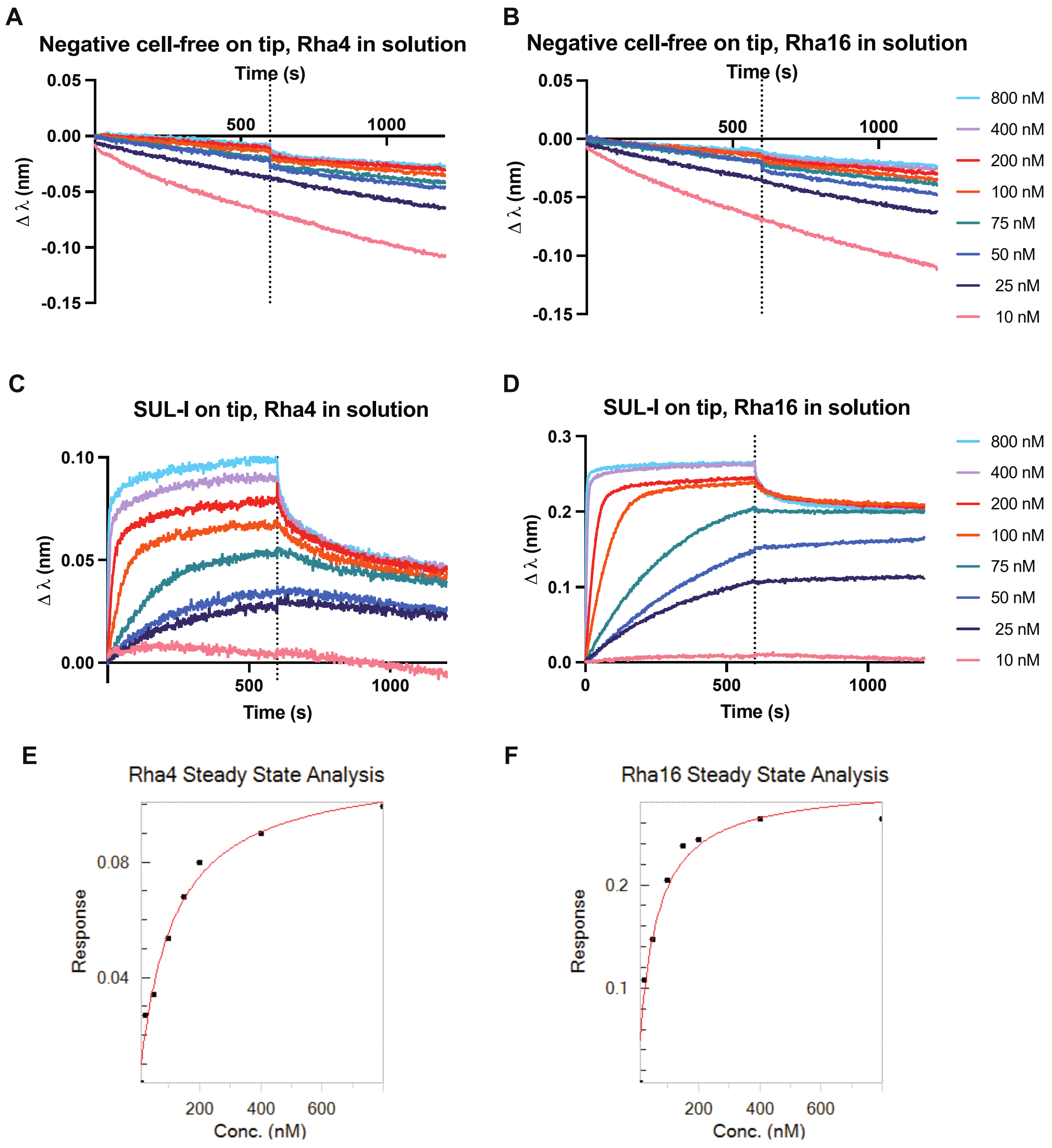


**Figure S8. Estimation of kinetic parameters with rhamnose substrate in solution and cell-free expressed lectin on tip.** Dilutions of (A) Rha4 or (B) Rha16 in solution with negative control cell-free reaction (no DNA) immobilized on the tip. No reference sensor was subtracted. Dilutions of (C) Rha4 or (D) Rha16 in solution with cell-free SUL-I immobilized on the tip. The “no DNA” sensor was subtracted as the reference sensor. Steady state analysis performed in ForteBio DataAnalysis9 with the equation Response= $R_{max}C$ /($k_D$+C) where C is concentration of the rhamnose ligand in solution for (E) Rha4 and (F) Rha16 ligands. $k_D$ values are reported in **Table S4**.

### Supplementary Tables

**Table S1. DNA sequences used in this study.** DNA (without signal sequence in the case of SUL-I) was codon optimized for expression in *E. coli* and purchased in the pJL1 vector (Addgene # 69496) for cell-free expression.

| Construct | PDB | Description | DNA Coding Sequence |
|---|---|---|---|
| pJL1-SUL-I | 5H4S | Amino acids 25-308 of SUL-I with an N-terminal 6xHis-tag followed by a TEV site in the pJL1 vector | ATGGGTAGCAGCCATCACCATCATCATCATAGCAGCGGTGAAA<br>ATCTGTATTTTCAGAGCTCCATGGGCGCAGTTGGTCGTACCTG<br>TGAAGGTAAAAGCCTGGATCTGGAATGTCCGGAAGGTTATATC<br>ATTAGCGTGAACTATGCAAACTATGGTCGTAATAGTCCGGGTA<br>TTTGCCCGCATAAAAGCAGCAATGCACCGCCTTGTAGCGCAAG<br>CAGCAGCCTGCGTATTATCAATGAACATTGCGATGGTCGTAGC<br>AGCTGTAGCGTTCATGCAACCAATGATGTTTTTGGTGATCCGT<br>GTCGTGGTGTGTATAAATACCTGGAAGTTGATTATAGCTGCCG<br>TCGTGATCCGGATTGTCAGCGCGAACTGGATTGCGAAGGTAAT<br>AGCATTAATATGCTGTGTCCGTATGCAGAAACACCGGCAATTC<br>ATATTTGCTATGCCATGTATGGTCGTCAGACCAGCGAACCGGT<br>TTGTCCGAGCAAAAGCATTAGCACCACCAATTGTGCAGCAAGC<br>AGCTCACTGAGCACCGCACGTAGCGTTTGTGAAGGTCGTTCAG<br>AATGTAGCATTGCAGCAAGTAACGATGTGTTTGGCGATCCTTG<br>TATTGGCACCTACAAATATCTGGAAATCGATTACATTTGTGCA<br>CGTCGTGGTCGTTCATGCGAAGGTTCAAGTCTGACCCTGAGCT<br>GTAGCAGCGGTCAGACCATTAGCGTTCTGGATGCATTTTATGG<br>TCGCACCGCAGGTCCGGAAATTTGTAAAGGTAATGCACAGGAT<br>CAGAATTGCCGTGCCGAAAGCAGCCTGAATATTGTTCAGAGCG<br>CCTGTAATGGTCGCAGCTCATGCAGCGTTAATGCAAATAACAA<br>CGTGTTCGGTGATCCATGTGTTGGGACATATAAGTATCTGGAA<br>GTGCTGTATAAATGCGCCTAACTCGAG |
| pJL1-CSL3 | 2ZX2 | Amino acids 1-195 of CSL3 with an N-terminal 6xHis-tag followed by a TEV site in the pJL1 vector | ATGGGTAGCAGCCATCACCATCATCATCATAGCAGCGGTGAAA<br>ATCTGTATTTTCAGAGCTCCATGGGCGCGATCTCAATCACATG<br>CGAGGGATCTGACGCTCTTCTTCAATGTGACGGCGCTAAGATC<br>CACATAAAGCGAGCAAACTACGGACGGAGACAACACGACGTGT<br>GTTCAATCGGCCGCCCCGACAACCAATTAACAGACACAAACTG<br>TCTCTCTCAATCGAGTACATCTAAGATGGCGGAGCGATGCGGC<br>GGCAAGTCTGAGTGCATCGTACCAGCTAGTAACTTCGTCTTCG<br>GAGACCCCTGCGTGGGAACTTACAAGTATCTTGACACAAAGTA<br>CTCATGTGTACAACAACAGGAGACGATCTCTAGTATCATCTGT<br>GAGGGGTCAGACTCACAACTCTTATGTGACAGAGGCGAGATCA<br>GAATCCAACGCGCCAATTACGGCCGACGTCAGCATGACGTTTG<br>CTCCATCGGGCGCCCGCACCAACAACTTAAGAACACGAACTGT<br>CTTTCTCAAAGTACGACATCTAAGATGGCAGAACGTTGCGACG<br>GGAAGCGCCAATGCATAGTCTCCGTATCTAACAGTGTATTCGG<br>TGACCCATGTGTAGGCACCTATAAATACTTGGACGTGGCTTAC<br>ACTTGTGACTAACTCGAG |

**Table S2. IC50 values determined for sugar inhibitors to disrupt binding between SUL-I in solution and rhamnose monosaccharide immobilized on tips.** Average IC50 and standard deviation from two independent experiments. IC50 data are plotted in **Figure S5** and raw data for one replicate is shown. * Asterisk indicates IC50 estimated by PRISM fit, but inhibition was not validated in the concentration range tested.

| **Inhibitor** | **Average IC50 (mM)** | **Std Dev IC50 (mM)** |
|---|---|---|
| L-rhamnose | 0.07 | 0.01 |
| Gb3 | 0.15 | 0.004 |
| isoGb3 | 0.30 | 0.03 |
| D-lactose | 0.27 | 0.02 |
| D-galactose | 1.26 | 0.08 |
| D-mannose | *169.80 | 31.25 |
| D-glucose | *352.85 | 40.52 |
| L-fucose | *476.55 | 1.63 |

**Table S3. Kinetic parameters determined from immobilization of rhamnose substrate on the sensor tip.** The $k_{obs}$ values are plotted from a 1:1 association model fit as a function of SUL-I concentration and fit with a simple linear regression in Graphpad PRISM to determine the values shown in the table below. The relationship $k_{observed}= k_{on}*C + k_{off}$ was used to determine $k_{on}$ and $k_{off}$ from the linear fit. The dissociation constant $k_d$ was estimated using the relationship $k_d= k_{off}/k_{on}$. The data represent the average and standard deviation of two replicates. The data for one replicate is shown in **Figure S6.**

| **Substrate on tip** | **Average $k_{on}$ ($nM^{-1}$ $s^{-1}$)** | **Std Dev $k_{on}$ ($nM^{-1}$ $s^{-1}$)** | **Average $k_{off}$ (1/s)** | **Std dev $k_{off}$ (1/s)** | **Average $k_D$ (nM)** | **Std Dev $k_D$ (nM)** |
|---|---|---|---|---|---|---|
| monosaccharide | 3.48E-05 | 1.24E-06 | 2.87E-03 | 9.19E-06 | 82.59 | 3.21 |
| Rha4 | 3.13E-05 | 4.95E-08 | 1.16E-03 | 4.10E-05 | 37.04 | 1.37 |
| Rha16 | 1.92E-05 | 1.63E-07 | 6.36E-04 | 1.98E-05 | 33.03 | 0.75 |
| PAA Rha | 1.98E-05 | 7.07E-08 | 8.64E-04 | 1.72E-04 | 43.65 | 8.53 |

**Table S4. Kinetic parameters determined from cell-free lectin immobilized on the sensor tip.** The $k_d$ was calculated by performing steady state analysis in ForteBio Data Analysis 9.0 for each experimental set up (eight concentrations of rhamnose substrate in solution for each replicate). The data represents the average and standard deviation of two replicates. The data for one replicate is shown in **Figure S8**.

| **Substrate in solution** | **Average $k_D$ (nM)** | **Std Dev $k_D$ (nM)** |
|---|---|---|
| Rha4 | 115 | 21 |
| Rha16 | 48 | 3 |

### Supplementary Materials and Methods

#### Cell-free Protein Synthesis

Lysate Preparation

SHuffle T7 Express (NEB) cells were grown in shake flasks at the 1 L scale. Cells were inoculated at optical density at 600 nm (OD600) = 0.05 and grown in 2xYTPG at pH 7.2 supplemented with 60 $\mu$g/mL spectinomycin and 100 $\mu$g/mL streptomycin at 37 °C. Cells were induced at $OD_{600}$ = 0.6 with 0.5 mM of IPTG to express T7 RNA polymerase and harvested at $OD_{600}$ = 3. Unless otherwise stated, all subsequent steps were performed on ice. Cells were harvested by centrifugation at 5,000 x g for 15 minutes and then washed 3 times with S30 buffer (10 mM Tris acetate pH 8.2, 14 mM magnesium acetate and 60 mM potassium acetate). Following washing, cells were pelleted at 10,000 x g for 2 minutes, then flash frozen and stored at -80 °C. Cells were resuspended in 1 mL/g S30 buffer. Then, cells were homogenized using an EmulsiFlex-B15 homogenizer (Avestin, Inc. Ottawa, ON, Canada) with 1 pass at a pressure of ~ 21,000 psig. Following lysis, cells were centrifuged for 10 minutes at 12,000 x g. Supernatant was collected and centrifuged again for 10 minutes at 12,000 x g. Supernatant was flash frozen and stored at -80 °C as the final extract.

Cell-free Protein Synthesis Reactions

Cell-free protein synthesis reactions were run at the 200 $\mu$L scale in 24 well plates or at the 15 $\mu$L scale in 2 mL tubes. Reactions were run as described previously using a modified PANOx-SP (PEP) formulation (Warfel et al. 2022) with the following modifications for disulfide bond formation. T7 Shuffle extract was incubated with 50 $\mu$M IAM at room temperature for 30 min before reaction set up. Reactions were supplemented with 10 $\mu$M DsbC, 1 mM GSH, and 4 mM GSSG. Reactions were incubated at 16°C for 20 hours unless otherwise noted.

Lectin Quantification

10 $\mu$M $^{14}$C-Leucine was supplemented into cell-free reactions for lectin quantification as previously described (Warfel et al. 2022). Reactions were centrifuged at 16,000 xg for 10 min and the supernatant was considered the soluble fraction. An autoradiogram was run with the soluble fraction of the cell-free reactions to confirm full length expression of the radiolabeled lectins. The gel was exposed for 5 days and imaged using a Typhoon imager. Lectin yields were plotted using Graphpad PRISM.

Affinity Purification

Alpha-Lactose Separopore® 6B-CL (Epoxy-Coupled) resin (bioPLUS Chemicals) was used to purify SUL-I and Immobilized D-Galactose Resin (Thermo) was used to purify CSL3. A 1:1 ratio of resin bed volume: soluble cell-free reaction was used for purifications. Micro Bio-Spin Chromatography Columns from Bio-Rad (Cat # 732-6204) were used with centrifugation at 700 x g for 1 min for separation. Manufacturer's protocol was modified such that soluble cell-free reaction was diluted 1:1 in buffer 1 (0.1 M $KH_2PO_4$, 0.15 M NaCl at pH 7.2) to equilibrated columns. Columns were washed at least 6 times with 3 CV buffer 1 before collection of 3, 1x CV elution fractions with 0.1 M of D-lactose or D-galactose in buffer 1.

**Bio-layer interferometry (BLI)**

All BLI experiments were performed using an Octet RED96 Instrument with data collected with ForteBio DataAcquisition9, analyzed and fit with ForteBio DataAnalysis9, and plotted with Graphpad PRISM.

Ligands

Biotinylated rhamnose monosaccharide (L-Rha$\alpha$-sp3-biot) and PAA rhamnose (L-Rha$\alpha$-sp3-PAA-biot) were purchased from GlycoNZ. Tetravalent (Rha4) and hexadecavalent (Rha16) cyclopeptides functionalized with L-rhamnose by triazole linkages, and biotinylated versions, were synthesized as reported previously (Liet et al. 2019).

Assays with Streptavidin tips

*Immobilization:* Biotinylated rhamnose substrates were prepared at a concentration of 1 $\mu$M for immobilization on streptavidin tips. Tips were incubated in PBS for 10 minutes before the loading procedure which consisted of: (1) baseline (PBS, 60s), (2) loading (biotinylated rhamnose substrate, 400 s), and (3) equilibration (PBS, 180s). The loading was followed by the blocking sequence described below.

*Blocking:* All tips were "blocked" by incubation in a well with 1:50 blank cell free reaction diluted in PBS to allow for any non-specific binding interactions of the cell-free reaction components with the SA tip to occur. The blocking procedure consisted of: (1) baseline (PBS, 60s), (2) association (1:50 blank cell free reaction diluted into PBS, 300s), and (3) dissociation (PBS, 200 s).

*Reference sensors:* Reference sensors were not loaded with biotinylated ligand and only underwent the blocking process. Reference sensors were dipped into the same solutions as the tips with immobilized substrates.

*Interaction:* Cell free reactions were diluted in PBS at the specified concentration and loaded into a 96 well plate for a total of 180 $\mu$L per well. For each sample the interaction assay consisted of: (1) baseline (PBS, 60s), (2) association (cell-free reaction diluted in PBS at desired concentration, 600s), and (3) dissociation (PBS, 200s). Tips were regenerated in 1 M rhamnose solution between repeats of the interaction sequence with each well.

*Competition:* Cell-free reaction expressing lectin diluted 1:50 in PBS was incubated with competitor sugar at a given concentration in a total volume of 200 $\mu$L per well in a 96 well plate for one hour at room temperature before running the BLI assay and following the interaction protocol above. Tips were loaded with biotinylated monosaccharide as described in the immobilization and blocking section.

Assays with Ni-NTA tips

*Immobilization:* 180 $\mu$L of cell-free reaction containing the His-tagged lectin was prepared at a 10-fold dilution in PBS and added to the 96 well plate. Tips were regenerated with $Ni^{2+}$ according to manufacturer's protocol before loading. The loading sequence was as follows: (1) baseline (PBS, 60s), (2) loading (His-tagged lectin substrate in CFPS, 900s), and (3) equilibration (PBS,180s).

*Reference Sensors:* Reference sensors were prepared according to the immobilization protocol above except were loaded with a negative control cell-free reaction (no DNA) instead of one containing lectin to account for non-specifically bound material that was observed to dissociate from the tips over time. Reference sensors were dipped into the same solutions as tips with immobilized substrates.

*Interaction:* For each sample, the interaction procedure consisted of: (1) baseline (PBS, 60s), (2) association (rhamnose substrate in PBS in solution, 600s), (3) dissociation (PBS, 600s). For *Shigella flexneri 2a* oligosaccharide interaction, association step was increased to 3600s. Tips were regenerated in 1 M rhamnose solution between repeating the interaction sequence with each well.

Kinetic Analysis:

SA tips: To determine $k_{observed}$, association curves were fit with a 1:1 model in ForteBio Data Analysis 9.0. The $k_{observed}$ value was plotted as a function of the concentration (C) of the lectin in solution and fit with a simple linear regression model in Graphpad PRISM. The relationship $k_{observed}= k_{on}*C + k_{off}$ was used to determine $k_{on}$ and $k_{off}$ from the linear fit. The dissociation constant $k_d$ was estimated using the relationship $k_d= k_{off}/k_{on}$.

Ni-NTA tips: Steady state analysis was performed to estimate $k_d$ using ForteBio Data Analysis 9.0. Response was calculated with an average from 590.0 to 595.0 seconds during a 600s association phase.

IC50 Analysis:

Association curves were fit with a 1:1 model in ForteBio Data Analysis 9.0 to determine the equilibrium binding signal ($R_{eq}$). The $R_{eq}$ of the interaction without any inhibitor was fixed at 100 and the $R_{eq}$ for each binding interaction curve corresponding with different inhibitor concentrations was normalized to the $R_{eq}$ of the interaction without any inhibitor. For interactions where an $R_{eq}$ was not determined (response $\leq$0), it was considered that normalized $R_{eq}$=0. As previously described to analyze inhibition using BLI (Orthwein et al. 2021), the normalized response was plotted against the log of the competitor concentration and fit using a nonlinear fit of log(inhibitor) vs normalized response using Graphpad PRISM.
